# Image-based metric of invasiveness predicts response to adjuvant temozolomide for primary glioblastoma

**DOI:** 10.1101/509281

**Authors:** Susan Christine Massey, Haylye White, Paula Whitmire, Tatum Doyle, Sandra K. Johnston, Kyle W. Singleton, Pamela R. Jackson, Andrea Hawkins-Daarud, Bernard R. Bendok, Alyx B. Porter, Sujay Vora, Jann N. Sarkaria, Maciej M. Mrugala, Kristin R. Swanson

## Abstract

Temozolomide (TMZ) has been the standard-of-care chemotherapy for glioblastoma (GBM) patients for more than a decade. Despite this long time in use, significant questions remain regarding how best to optimize TMZ therapy for individual patients. Understanding the relationship between TMZ response and factors such as number of adjuvant TMZ cycles, patient age, patient sex, and image-based tumor features, might help predict which GBM patients would benefit most from TMZ, particularly for those whose tumors are not MGMT methylated. Using a cohort of 90 newly-diagnosed GBM patients treated according to the Stupp protocol, we examined the relationships between several patient and tumor characteristics and volumetric and survival outcomes during adjuvant chemotherapy. Volumetric changes in MR imaging abnormalities during adjuvant therapy were used to assess TMZ response. T1Gd volumetric response is associated with younger patient age, increased number of TMZ cycles, longer time to nadir volume, and decreased tumor invasiveness. Moreover, increased adjuvant TMZ cycles corresponded with improved volumetric response only among more nodular tumors, and this volumetric response was associated with improved survival outcomes. Finally, in a subcohort of patients with known MGMT methylation status, MGMT methylated tumors were more diffusely invasive than unmethylated tumors, suggesting that the improved response in nodular tumors is not driven by a preponderance of MGMT methylated tumors. Our finding that less diffusely invasive tumors are associated with greater volumetric response to TMZ suggests that patients with these tumors may benefit from additional cycles of adjuvant TMZ, even for those without MGMT methylation.

## Introduction

Glioblastoma (GBM) is the most common malignant primary brain tumor found in adults (1). Despite diligent research efforts, patients diagnosed with this aggressive cancer have a median overall survival of 15 months (2,3). In 2005, Stupp et al. found that maximal safe resection followed by concurrent radiotherapy and temozolomide (TMZ) chemotherapy and six adjuvant cycles of TMZ resulted in a median overall survival of 14.6 months compared to 12.1 months for radiotherapy alone. Today, this protocol remains the standard-of-care for patients diagnosed with GBM.

As an alkylating agent, TMZ operates by methylating and damaging DNA, preventing proliferation and inducing apoptosis (4). Compared to other therapeutic agents, TMZ is relatively blood-brain barrier (BBB) penetrant, with a CSF to plasma ratio of 33% (5), which is one factor that makes it effective against gliomas. The angiogenic nature of glioblastoma causes the breakdown of the BBB in the vicinity of the tumor, which also contributes to the drug’s ability to reach the tumor cells (6,7). In addition to inducing the apoptosis of glioma cells, TMZ in combination with radiotherapy can cause pseudoprogression, which is observed as progressive imaging changes that look similar to true progression and are thought to be a result of treatment-induced inflammation (7). The similar radiological presentation of growing tumor and pseudoprogression complicates the assessment of TMZ response (8). Some reports suggest that waiting until after three cycles of adjuvant TMZ (i.e., approximately 12 weeks from completion of radiotherapy) to assess treatment response can improve the accuracy of progression determination (7).

Patients typically receive a daily TMZ dose of 75 mg per square meter of body-surface area during radiotherapy, followed by a dose of 150-200 mg per square meter for 5 days during each 28 day adjuvant cycle for 6-12 cycles. TMZ is generally well tolerated, with about one-third of patients experiencing nausea and vomiting that is typically well controlled by anti-emetics (7). Patients are also at risk for infection, lymphopenia, neurotoxicity (7), or hematologic toxicities, such as thrombocytopenia (8). Stupp et al. found that in a population of over 200 patients, the percentage of people who discontinued therapy due to the toxic effects of TMZ was only 5% during the concurrent stage and 8% of patients during the adjuvant stage (2). The FDA labeling specifies giving six cycles of adjuvant TMZ, although the number of cycles of adjuvant TMZ administered in clinical practice varies. Administration of the drug may be discontinued early due to adverse effects or disease progression, while some patients and their physicians elect to administer the drug beyond 6-12 cycles, sometimes for as long as 2-3 years or until ultimate disease progression (8). The relationship between number of adjuvant cycles received and outcome has not been clearly elucidated. Three studies found that patients who received more than 6 cycles of TMZ had improved survival compared to those who received less (9–11), while another study found no survival difference between these two groups of patients (12).

Even when patients receive the same number of cycles of TMZ without adverse effect, there can be large variation in tumor response. This is largely attributed to particular molecular features (genetic and epigenetic), which may predispose a patient to a better TMZ response and/or delayed evolution of TMZ resistance. The molecular feature that is given the most attention in regards to TMZ sensitivity is O6-methylguanine-DNA methyltransferase (MGMT) promoter methylation (8). Methylation of the MGMT promoter in GBM effectively silences this DNA repair gene, making tumor cells unable to repair the cytotoxic O6-methylguanine lesions induced by TMZ and some other alkylating agents (13–15). MGMT promoter methylation exists in about 35% of GBMs (16) and is associated with longer overall and progression-free survival (14,15). Research has suggested that tumor responsiveness to TMZ is also impacted by IDH1 mutation and p53 mutation in lower grade gliomas (17,18).

While genetic differences are currently the best supported predictors of TMZ response, recent studies have found that other patient characteristics impact TMZ response. Recently, a large-scale investigation found that female GBM patients live longer than male GBM patients (19). Considering that TMZ is a part of standard-of-care practice, this raises the question of whether there is an impactful sex difference in tumor responsiveness to TMZ. It has been observed that females have an improved volumetric response and exhibit better tumor control during adjuvant TMZ than males (20,21), but further research is needed to fully elucidate the biological mechanism of this sex difference. Age is recognized as a significant prognostic indicator for GBM patients. It is further thought that older patients are not as tolerant to aggressive treatment as their younger counterparts (22), and patients older than 70 years were not included in the study that established the current standard-of-care (2). While toxicity and adverse reactions remain a concern, a prospective study on GBM patients 65 years or older found that adding adjuvant TMZ to a radiotherapy treatment course improves median overall survival by 3.7 months and PFS by 5.4 months (23). This substantial impact on outcome emphasizes the need to assess whether age impacts tumor responsiveness to TMZ.

Considering the large variation in response to TMZ, the potential for adverse reaction, and the uncertainty caused by pseudoprogression, deciding how many cycles of adjuvant TMZ to administer to a patient is a challenging task for clinicians. The possibilities of adding another therapy during adjuvant TMZ, such as tumor treating fields (TTF), or continuing administration of TMZ beyond six cycles adds further complexity to clinical decision making. Outside of the presence of MGMT methylation, which does not apply to a majority of patients, there are few clear indicators to aid clinicians in this process. In this investigation, we sought to identify image-based characteristics associated with TMZ response that can be assessed in the pre-adjuvant setting. By comparing pre-adjuvant and post-adjuvant MR images, we sought characteristics that are associated with volumetric response, overall survival, and progression free survival. Additionally, we examined how the number of TMZ cycles received and MGMT methylation status influenced these relationships.

## Methods

### Patient Cohort

Our lab has amassed a multi-institutional repository of over 1400 glioma patients. The cohort for this present study consists of all patients in this repository who met the following criteria: A) diagnosed with primary GBM (n=1323), B) received maximal safe resection, concurrent radiation therapy (XRT) and TMZ, and at least one adjuvant cycle of TMZ (n=234), C) had available age at diagnosis, sex, overall survival, and treatment start/stop dates (n=210), D) did not receive any therapies other than XRT, TMZ, anti-seizure medications, or steroids between the first surgery and first cycle of adjuvant TMZ (n=175), and E) had sufficient pre-adjuvant and post-adjuvant MR imaging (detailed below in “Imaging and Biomathematical Model”) (n=90). These inclusion criteria resulted in the identification of a cohort of 90 patients **(Table 1)**. Eleven patients received a therapy other than TMZ concurrent with or in between cycles of TMZ; these other therapies included additional resection or radiotherapy, thalidomide, accutane, and bevacizumab. While the concurrent use of TMZ and TTF is becoming more common, none of the patients in this cohort received TTF during adjuvant therapy. Further, to ensure that we captured the effect of TMZ exclusively in these eleven cases, the image before the start of the other therapy was used as the post-adjuvant image, so that no patients received other therapies during the analyzed imaging period.

**Table 1:** Distributions and counts of relevant demographic, volumetric, and treatment-based patient characteristics. Extent of resection is abstracted from surgical notes and radiological reports and is not uniformly determined radiographically. Distributions of the nadir-related variables are in **Supplement 1.** (LTFU = lost to follow-up)

|  | N= | Mean | Median | Range |
| --- | --- | --- | --- | --- |
| Sex |  |  |  |  |
| Male | 60 (66.7%) | ---- | ---- | ---- |
| Female | 30 (33.3%) | ---- | ---- | ---- |
| Age (years) | 90 | 54.66 | 57.5 | 18-76 |
| Overall Survival (days) |  |  |  |  |
| Confirmed death | 71 (78.9%) | 806.0 | 562 | 115-3245 |
| Alive/LTFU | 19 (21.1%) | 1404 | 1278 | 128-3819 |
| Time from adjuvant TMZ to progression (days) | 43 (47.8%) | 241.0 | 30 | 7-1709 |
| Extent of Resection |  |  |  |  |
| Gross Total Resection | 40 (44.4%) | ---- | ---- | ---- |
| Sub-total Resection | 35 (38.9%) | ---- | ---- | ---- |
| Biopsy | 15 (16.7%) | ---- | ---- | ---- |
| Cycles of adjuvant TMZ<sup>a</sup> | 90 | 6.122 | 5 | 1-21 |
| Received <6 cycles | 47 (52.2%) | ---- | ---- | ---- |
| Received 6 cycles | 14 (15.6%) | ---- | ---- | ---- |
| Received 7+ cycles | 29 (32.2%) | ---- | ---- | ---- |
| Pre-adjuvant D/rho (mm<sup>2</sup>) | 90 | 2.073 | 1.409 | 0.0034-9.525 |
| Pre-adjuvant T1Gd radius (mm) | 90 | 12.03 | 10.81 | 2.312-32.25 |
| Post-adjuvant T1Gd radius (mm) | 90 | 11.90 | 11.62 | 0.00-22.22 |
| %Δ T1Gd | 90 | 7.70% | -0.16% | -100% - 260% |

<sup>a</sup>The cycles of adjuvant TMZ reported here exclude any cycles that were given in conjunction with other anti-tumor therapies since these were excluded from our analysis (see **Methods**). It should be noted that the majority of patients did receive at least 6 cycles of TMZ, even if they were not counted for the adjuvant period in our analysis.

### Imaging and Biomathematical Model

We defined “adjuvant TMZ” as the time period when patients consistently received cycles of TMZ alone after the completion of surgery and concurrent TMZ and XRT. If patients received another therapy concurrent with TMZ or between cycles of TMZ, we only considered the period when they received TMZ alone to be “adjuvant TMZ” and excluded the cycles administered after the start of the other therapy. Further, in order for patients to be included in this study, they had to have the following available MR images: 1) gadolinium enhanced T1-weighted (T1Gd) and T2-weighted or T2 fluid-attenuated inversion recovery (T2-FLAIR) images between concurrent XRT/TMZ and adjuvant TMZ (“pre-adjuvant” images), and 2) T1Gd and T2-FLAIR images near the dates of their last cycle of adjuvant TMZ (taken either 1-2 cycles before the end of adjuvant therapy or up to 40 days after) (“post-adjuvant” images). All post-adjuvant images were taken after the administration of cycles of TMZ only and before the start of any other therapy. The relevant abnormality in these images were segmented by trained individuals with the assistance of our in-house thresholding-based software to calculate a three-dimensional tumor volume. Each tumor volume was then used to determine a spherically equivalent tumor radius for use in this investigation (T1Gd radius and T2-FLAIR radius). From the T1Gd radii we computed the percent change of the T1Gd radius over the adjuvant TMZ cycles (%Δ T1Gd), which is calculated by finding the difference between the post-adjuvant T1Gd radius and pre-adjuvant T1Gd radius and dividing it by the pre-adjuvant T1Gd radius.

Next, using the pre-adjuvant T1Gd and T2-FLAIR images, we calculated a mathematical model–based tumor invasion metric called D/rho. This metric derives from the proliferation-invasion (PI) model of glioblastoma growth (24,25) and describes the ratio of overall tumor invasion to proliferation, with higher D/rho indicating a more diffuse tumor and lower D/rho indicating a more nodular tumor (26,27). Diffuse tumors have higher levels of model-predicted net cellular invasion relative to model-predicted net cellular proliferation, while nodular tumors have more proliferation relative to invasion into the surrounding tissues. We computed this metric at the pre-adjuvant imaging time point to establish a baseline of tumor invasiveness prior to the administration of adjuvant TMZ.

### Response Indicator

For the purposes of this investigation, we split our cohort into “responders” and “non-responders” based on the tumor volume changes observed in T1Gd images over the course of adjuvant therapy. Patients that had a decrease in T1Gd abnormality volume (negative %Δ T1Gd) following adjuvant TMZ were considered “responders” and patients that had an increase in volume (positive %Δ T1Gd) were considered “non-responders”. Note that this classification is not intended for clinical decision-making or to distinguish progression from stable disease; therefore, in this investigation, these terms refer exclusively to T1Gd volumetric response and “outcome” refers to survival. While we compared pre-adjuvant T1Gd volume with post-adjuvant T1Gd volume for the calculation of %Δ T1Gd, we also conducted an investigation that compared the pre-adjuvant volume with the nadir volume (%ΔT1Gd-Nadir). This nadir volume is defined as the smallest T1Gd volume at any time point after the pre-adjuvant time point and before or at (if a smaller volume was not reached earlier or no intermediate images were available) the post-adjuvant time point. Note that pseudoprogression, when it occurs, can increase our imaging-based measure of tumor volume. This could result in some tumors being misclassified as responders due to the resolution of pseudoprogression during the course of adjuvant TMZ. While this remains a potential confounder for many GBM studies, our classification correlated well with overall and progression free survival, suggesting that any such misclassification was minimal. To further reduce the possibility of misclassification due to pseudoprogression, we re-performed all of our analyses in a supplemental investigation using a subcohort of patients with more than 12 weeks between the end date of XRT and date of post-adjuvant imaging (n=72).

### Statistical Analysis

Two-sided t-tests with Welch’s corrections were used to test for differences in the means of two groups. F-tests with linear regression models were used to test whether two variables had a significantly positive or negative correlative relationship or were not related. Kaplan-Meier curves were used to visualize survival data and log-rank tests were used to test whether two groups had significantly different outcomes. All of these statistical tests were performed using R (28,29) using packages *survival* (30), *survminer* (31), *and ggplot2* (32). A p-value of 0.05 was used as the cut-off for statistical significance.

### Study Approval

All patients included in this investigation were consented prospectively or approved for retrospective research by institutional review boards.

## Results

### T1Gd volumetric response correlates with younger patient age, increased number of TMZ cycles, longer time to nadir, and decreased tumor invasiveness

In order to understand whether various patient or tumor characteristics were significant predictors of tumor response to TMZ, we classified patients as “responders” or “non-responders” based on change in T1Gd volume over the course of adjuvant TMZ, as detailed in the Methods. Responders (n=45) were younger (t-test, p=0.0450), received more cycles of TMZ (p<0.0001), reached nadir later during the adjuvant time period (p=0.0046), and had tumors that were more nodular (p=0.0191) than non-responders (n=45) **(Figure 1)**. MR images of a nodular responding patient and a diffuse non-responding patient are shown as examples in **Figure 2**. There was no difference in the pre-adjuvant tumor volume (T1Gd radius p=0.1007, T2-FLAIR radius p=0.719) between responders and non-responders. Responders also had significantly longer survival than non-responders (log-rank test, p=0.0028) **(Figure 3)**. Among patients with a recorded date of progression, responders (n=21) tended to have a longer time between TMZ and progression than non-responders (n=22) (p=0.0674) **(Figure 3)**. The relationship between the T1Gd-based volumetric response and outcome validates its relevance as an indicator of treatment response. Meanwhile, changes in T2-FLAIR abnormality had no clear relationship with patient outcome. Therefore, quantifying the changes in T2-FLAIR radius does not appear to provide a valuable response indicator.

**Figure 1:**
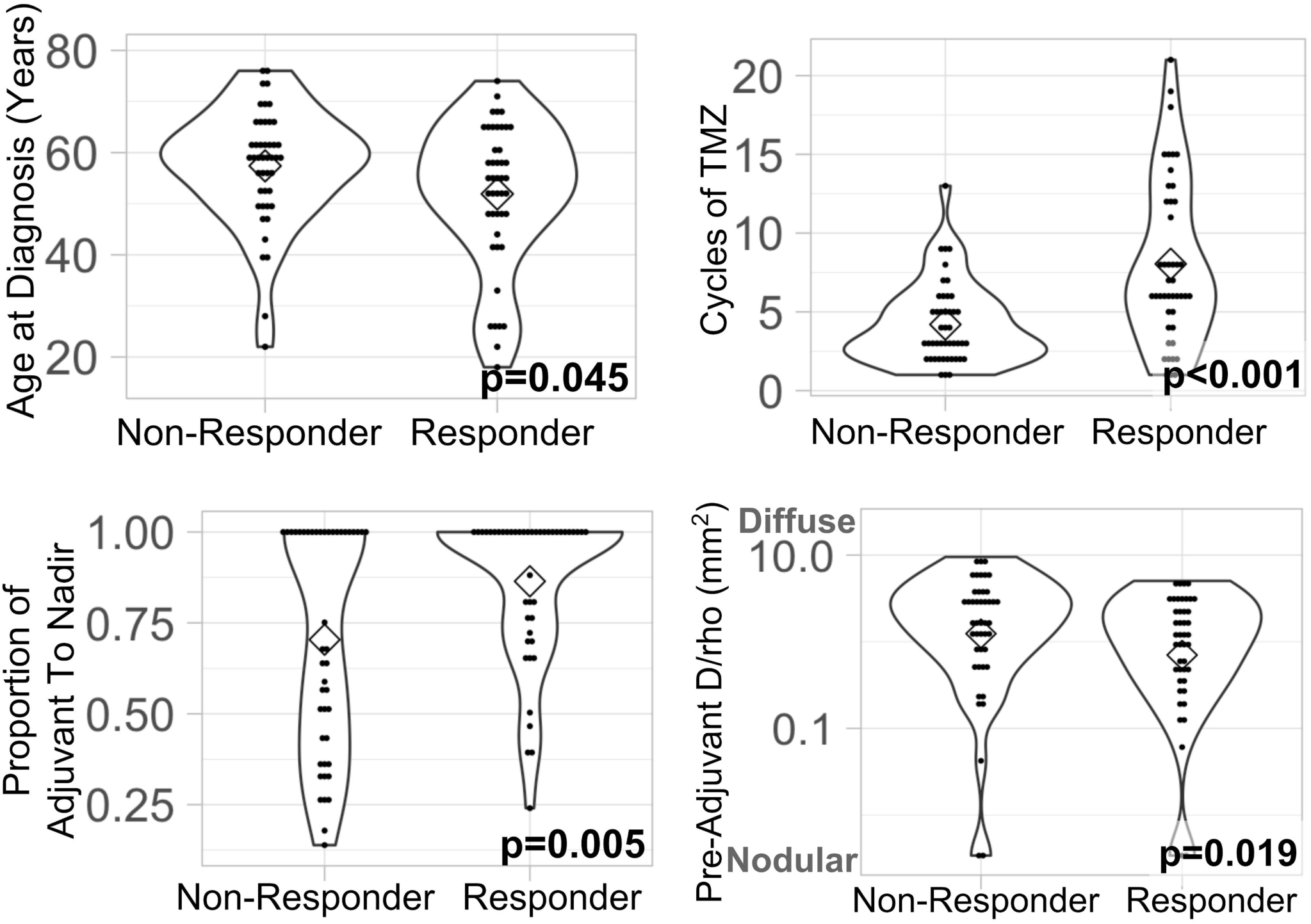
Characteristic differences between responders (n=45) and non-responders (n=45). Statistical tests (t-tests) show that volumetric responders (decrease in T1Gd volume during adjuvant TMZ) were younger, received more cycles of TMZ, reached nadir relatively later during adjuvant therapy, and had more nodular tumors than non-responders (increase in T1Gd volume during adjuvant TMZ). Proportion of adjuvant to nadir is calculated as the number of days between pre-adjuvant and nadir images divided by the total number of days between pre-adjuvant and post-adjuvant images.

**Figure 2:**
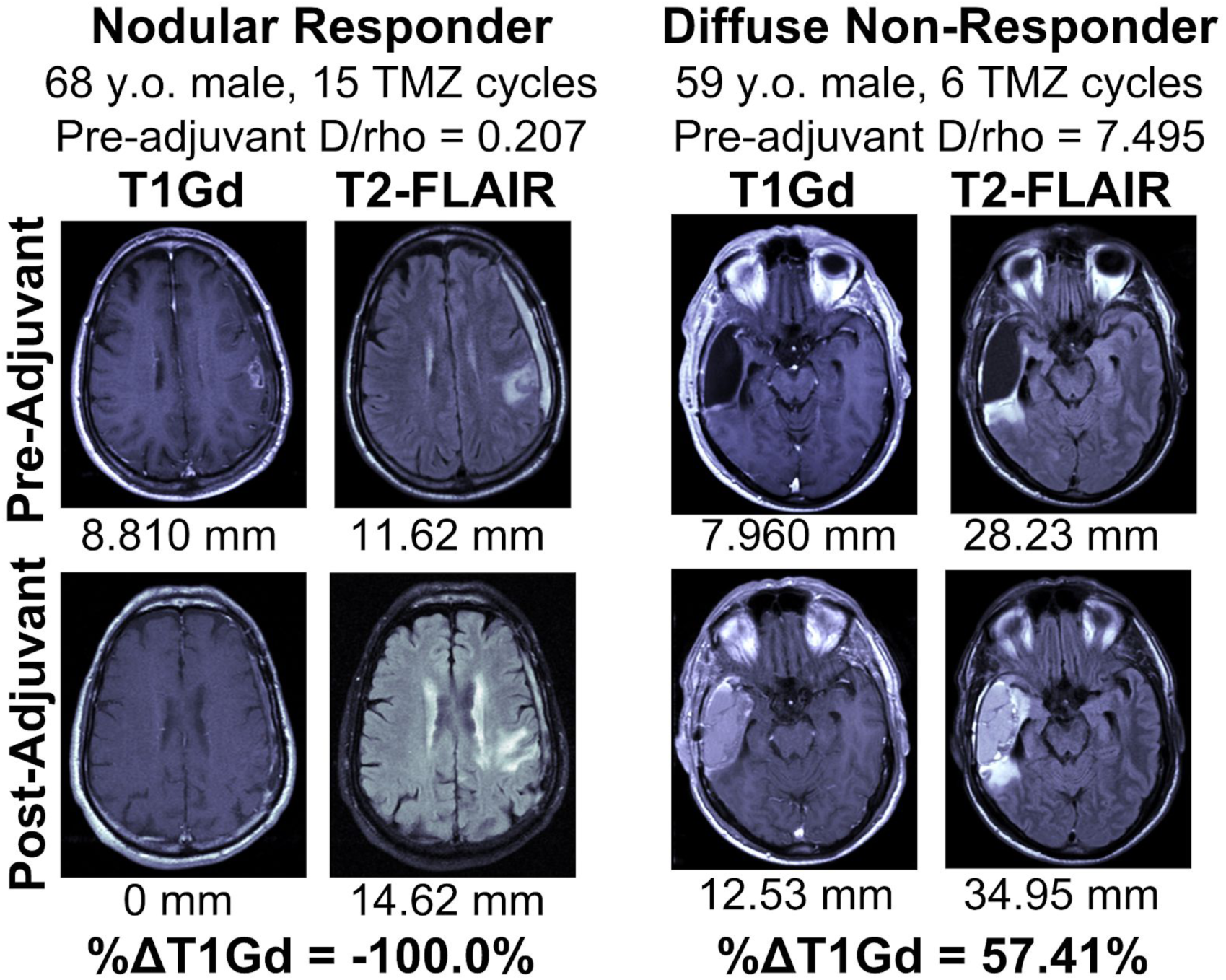
Pre-adjuvant and post-adjuvant T1Gd and T2-FLAIR MR images of a nodular responding patient and a diffuse non-responding patient. The spherically-equivalent radius converted from the volume of each lesion is listed below the image in millimeters; these were used to derive the D/rho diffusivity index. (y.o. = years old)

**Figure 3:**
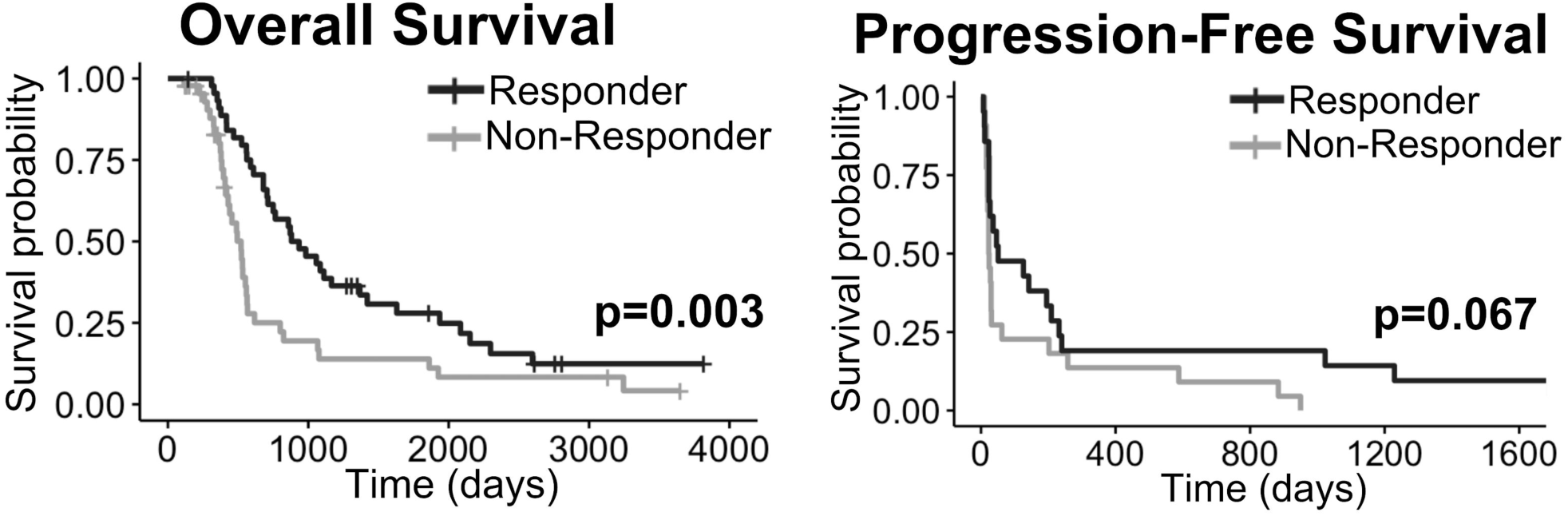
Responders (n=45, decrease in T1Gd volume during adjuvant TMZ) had significantly longer overall survival than non-responders (n=45, increase in T1Gd volume during adjuvant TMZ). Among patients with dates of progression, responders (n=21) tended to have longer times to progression than non-responders (n=22).

### Increasing number of cycles correlates with volumetric response only in nodular tumors

In order to test the impact of tumor invasiveness on volumetric response and outcome, we divided the patients into three equally sized groups based on pre-adjuvant D/rho (nodular, moderate, and diffuse). When we considered the impact of number of cycles on volumetric response, we found a significant negative correlation between cycles of TMZ received by a patient and their %Δ T1Gd among nodular tumors (F-test, p=0.0062), but this relationship did not exist among diffuse tumors (p=0.4040) **(Figure 4)**. This indicates that additional cycles of TMZ have a clearer volume reduction benefit among patients with nodular tumors than among those with diffuse ones. This volumetric benefit is also tied to outcome, with nodular tumors having a distinct relationship between volumetric change and survival. Specifically, we found that the survival difference observed between responders and non-responders is only significant among the nodular tumors (log-rank, nodular p=0.0021, diffuse p=0.793) **(Figure 4)**.

**Figure 4:**
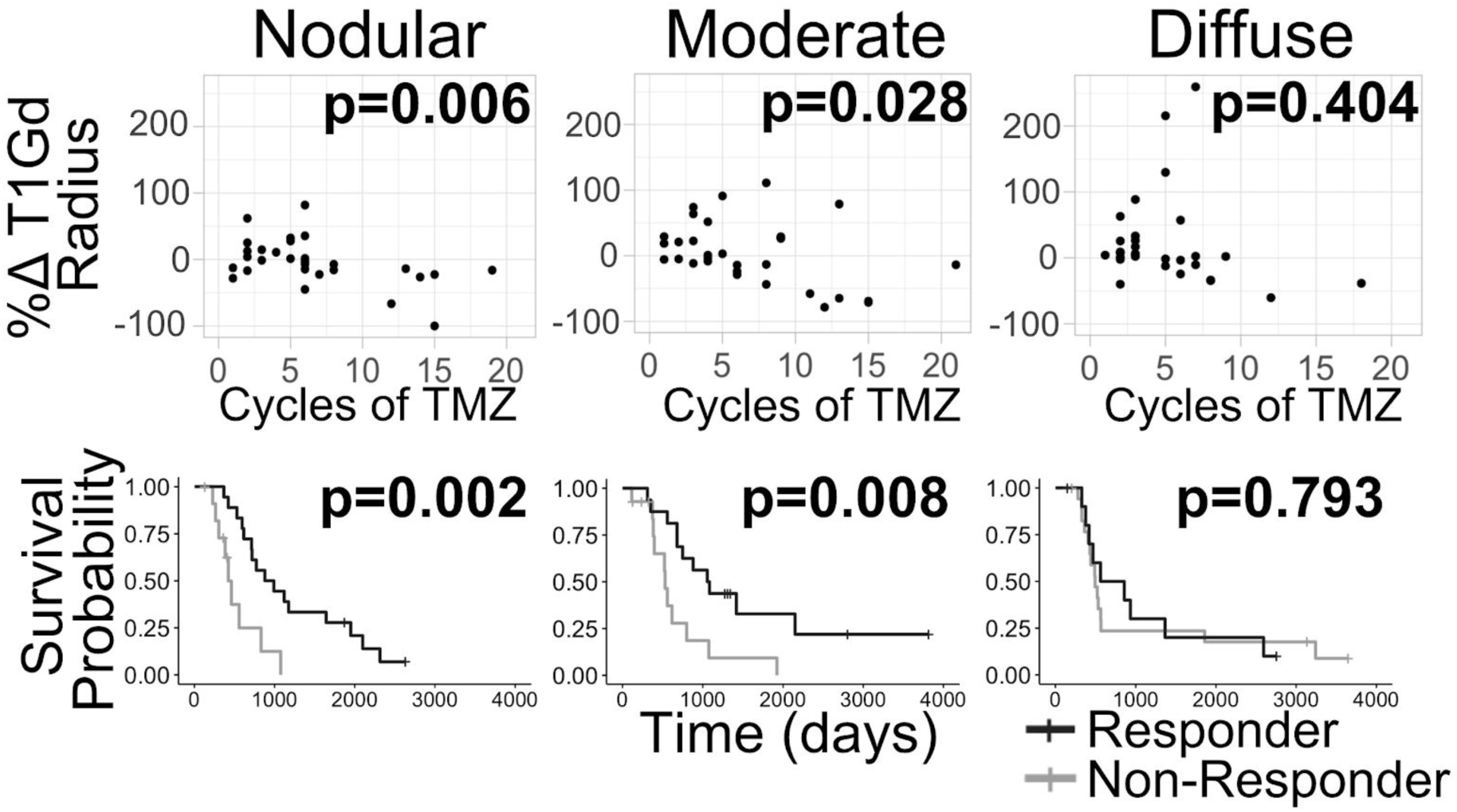
Patients were split into three evenly sized groups based on their pre-adjuvant D/rho (nodular, moderate, and diffuse). Among the nodular tumors (n=30), there is a significant negative correlation between volumetric response and cycles of TMZ received. Then this improved response is clearly tied to outcome since nodular responders (based on T1Gd volume change) had significantly longer survival than non-responders of the same group. The relationships between cycles, response, and outcome are not significant among diffuse tumors (n=30).

### MGMT methylated tumors are more diffusely invasive

Since methylation of the MGMT promoter corresponds with improved TMZ response, we investigated whether our findings might simply be attributable to a co-occurrence of those features with MGMT methylation. Using our limited sample of patients with known MGMT methylation status (methylated n=9, unmethylated n=14), we analyzed the relationship between methylation status, tumor volumetric response, cycles of TMZ, and D/rho **(Supplement 2)**. Patients with MGMT methylated tumors had significantly better survival than those with unmethylated tumors (log-rank, p=0.014), consistent with existing literature. Further, those with MGMT methylated tumors are more commonly responders (6 responders vs. 3 non-responders) and have significantly better volumetric response than those with unmethylated tumors (t-test, p=0.024). Among the volumetric responders (n=11), MGMT methylation (n=6) showed a survival benefit over unmethylation (n=5) (p=0.014). However, patients with methylated tumors also received more cycles of TMZ (p=0.0156), which could indicate that prescribing practices have created a confounding factor in the relationship between methylation and volumetric response. Focusing within methylated tumors, we observe a clear negative correlation between cycles of TMZ received and %Δ T1Gd during adjuvant TMZ **(Figure 5)**, similar to that among nodular tumors. This comparison is limited by its small sample size, but supports the existing idea that methylated tumors respond well to TMZ chemotherapy. Interestingly, MGMT methylated tumors are more diffuse than unmethylated tumors (p=0.011). Among only unmethylated tumors, we again see the pattern that responders tend to have tumors that are more nodular.

**Figure 5:**
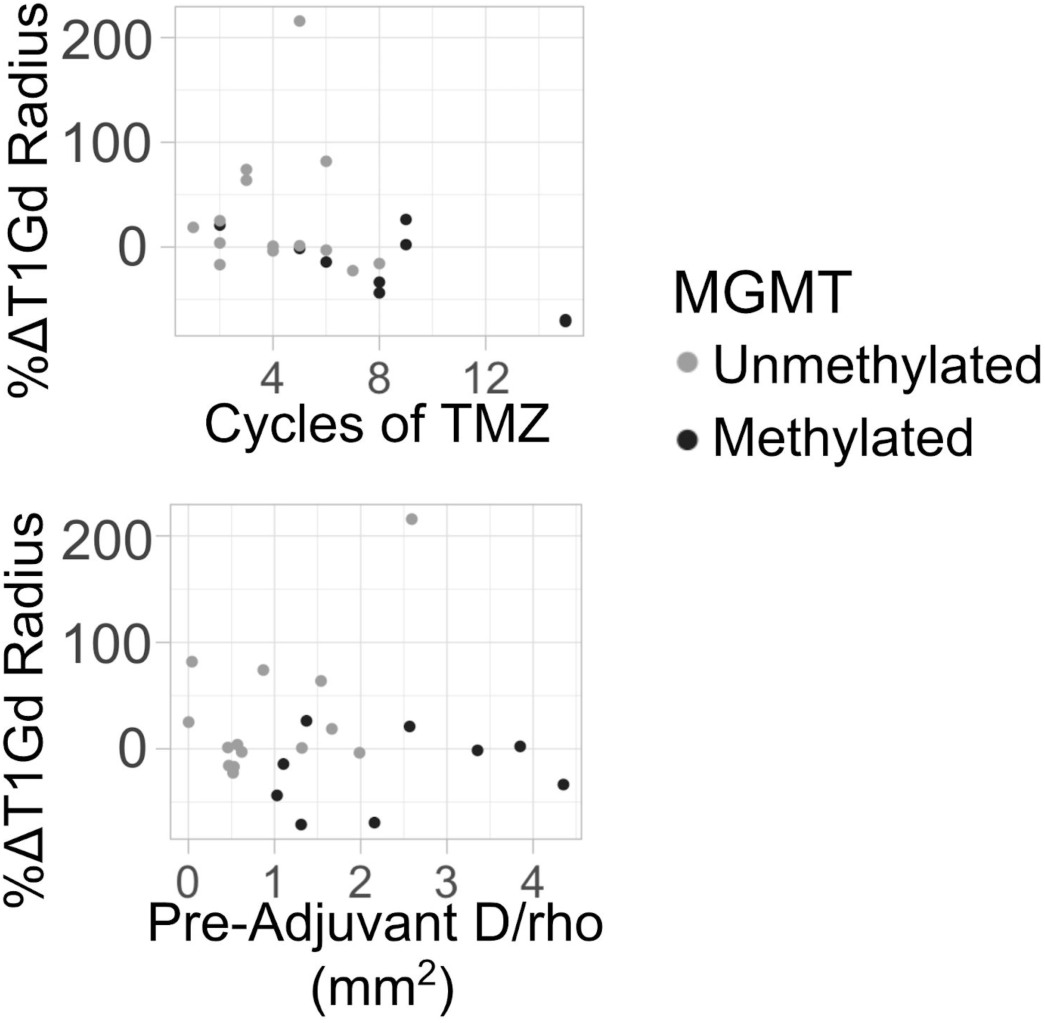
Methylated patients (n=9) have a clear negative trend between cycles of TMZ and volumetric response, while unmethylated patients (n=14) show a similar trend, but with more deviance. Methylated patients have more diffuse tumors (higher pre-adjuvant D/rho) than unmethylated.

Since our comparison of MGMT status and pre-adjuvant D/rho had a relatively small sample size, we identified 49 additional first-diagnosis GBM patients (who were excluded from other analyses because they did not meet the post-adjuvant imaging inclusion criteria) with available MGMT status and pre-adjuvant D/rho from our database for validation. In this combined cohort (23 patients who met inclusion criteria plus the 49 additional patients for this particular analysis), MGMT methylated patients (n=28) had tumors that were significantly more diffuse than those in unmethylated patients (n=44) (p=0.006) **(Supplement 3)**. This confirms that the relationship between nodularity and response is not confounded by a predominance of methylated tumors in the nodular group.

### Using volume change until nadir validates previous results

We investigated whether analyzing the nadir (lowest T1Gd volume during adjuvant therapy) time point instead of the post-adjuvant time point would be more informative **(Supplement 1)**. Using volume change between the pre-adjuvant T1Gd image and the nadir image for calculating the percent change in T1Gd radius (%Δ T1Gd-Nadir), we found the similar results to those shown in **Figure 4**. Specifically, among nodular tumors there was a significant negative correlation between number of TMZ cycles received and %Δ T1Gd-Nadir (F-test, p<0.0001), and this relationship was not significant among diffuse tumors (p=0.1610) **(Supplement 1)**. We also found that the overall volumetric change (from the pre-adjuvant to post-adjuvant time points) was more closely tied to clinical outcome than the change from pre-adjuvant imaging to nadir.

In order to assess the potential impact of pseudoprogression on our results, we performed all of the above analyses on a subcohort of patients that had at least 12 weeks between the XRT end date and the date of post-adjuvant imaging (n=72) **(Supplement 4)**. This investigation showed comparable results with the full cohort, with the sole exception being the comparison of pre-adjuvant D/rho between responders and non-responders, which only trended towards significance (p=0.0658).

## Discussion

In this investigation, we examined a number of patient attributes to assess whether any might be predictive of response to adjuvant TMZ. We found that patients whose T1Gd abnormality decreased in volume during adjuvant TMZ therapy were younger in age, received more cycles of TMZ, had longer time to nadir, and had more nodular tumors than those whose abnormality increased in volume. This decrease in volume was associated with better outcomes, including longer overall survival and a trend towards longer time to progression compared to those that had an increase in volume.

Some of these findings were expected and consistent with earlier studies. For example, younger patients have been shown to have better outcomes in other studies (22). While this could be caused by differences in chemotherapy tolerance, a more favorable volumetric response to chemotherapy could also contribute to the survival differences observed between older and younger GBM patients. Additionally, while toxicity remains a concern, our finding that increased cycles of TMZ correlates with volumetric response supports other studies showing that more cycles of TMZ result in better response and outcomes (9–11).

Other findings were less intuitive, but also consistent with earlier studies. The association of longer time to nadir with response to TMZ, while not expected, is consistent with longer durability of TMZ effect upon tumor. Initial pseudoprogression may also contribute to the observation that volumetric responders reached nadir volume later in their adjuvant cycling. Our analyses that used volume change between pre-adjuvant and nadir images supported the results from the analyses that used the change between pre-adjuvant and post-adjuvant volumes.

The most important and unanticipated finding of this work was that nodular tumors tend to respond more favorably to adjuvant TMZ, both in terms of volumetric change and outcomes. Patients whose T1Gd abnormality decreased in size over the course of adjuvant therapy had significantly more nodular tumors than those who had an increase in size. Furthermore, patients with more nodular tumors had a clear negative correlation between cycles of TMZ received and volumetric response, with more cycles of TMZ resulting in a more favorable volumetric response. Among diffuse tumors, this relationship was neither visibly clear nor statistically significant. When looking at clinical outcomes, volumetric responders had significantly longer overall survival compared to volumetric non-responders among patients with nodular tumors, while this comparison was not significant among patients with diffuse tumors.

It has been previously suggested that TMZ might be less effective in more diffuse tumors. One study suggested that TMZ might be present in higher concentrations near the contrast-enhancing core of the tumor, where the BBB is more likely to be compromised, compared to the surrounding tissue (7,33). In a nodular tumor, a larger proportion of the visible tumor cells (on MR imaging) are near the contrast-enhancing core of the tumor, while in diffuse tumors, there are thought to be more image-detected invasive cells in the periphery (34), potentially limiting the efficacy of TMZ. We think this is the most likely explanation for the observations we made in this investigation, but more research is needed to fully understand how tumor characteristics interact with the BBB to affect drug distribution in brain tissue.

Lack of MGMT expression is mechanistically linked to TMZ sensitivity, and MGMT promoter methylation results in more favorable responses to TMZ chemotherapy (8,13–15,22). Although limited by the small sample size of our patient cohort with known MGMT status, we wanted to ensure that the relationship between nodularity and responsiveness to TMZ was not confounded by MGMT methylation. We found that the MGMT methylated tumors were more diffuse at the pre-adjuvant imaging time point than the unmethylated tumors (**Supplement 2**). When we expanded our cohort to include more than seventy patients with MGMT status, we found that this observation remained true (**Supplement 3**). Therefore, we concluded that since the presumably more responsive MGMT methylated tumors were concentrated in the diffuse group, the observation that nodular tumors respond better to TMZ is not likely confounded by this molecular marker.

### Limitations and Future Work

We acknowledge that our retrospective investigation has some limitations and hope that after independent replication, these results will be validated and become clinically applicable. Further, our response classification (responder vs non-responder) is not intended for clinical use and is not meant to distinguish progression or stable disease or to be used for clinical decision-making. We focused on quantifiable imageable response to the exclusion of nuanced clinical aspects of patient response, such as performance status and steroid use, that are needed in clinical metrics like RANO. Despite the simplicity of our metric and the potential for pseudoprogression to confound its results, it remained closely tied to outcome, which we believe justifies its use in a retrospective analysis. While some patients did receive other therapies during their adjuvant TMZ, this only occurred in a small number of cases and usually towards the end of adjuvant therapy. Further, no patients received other therapies during the analyzed imaging periods.

Future work could attempt to identify other tumor characteristics that correspond to TMZ response. Notably, our investigation did not find a relationship between changes in the T2-FLAIR abnormality during adjuvant TMZ and tumor characteristics or patient outcome. While T2-FLAIR identifies fluid, this could be associated with extracellular fluid from leaky vasculature, immune recruitment and inflammation, or perhaps some other process. Each of these has different biological implications and more research is needed to uncover T2-FLAIR image features that indicate which of these processes are being visualized and to explore the different clinical implications of these processes. Future work could also look for sex differences in TMZ response. The results of previous work on sex differences suggests that TMZ might have sex-specific effects (20,21), which we hypothesized might affect the tumors in this cohort. When we ran the tests from this investigation on male and female patients separately, we observed that the same trends remained significant in the male cohort and were mostly insignificant in the female cohort **(Supplement 5)**. However, the small size of our female sample limits our ability to draw conclusions from this observation.

## Conclusion

In our retrospective investigation, we found that factors like patient age, cycles of TMZ received, time to nadir volume, and tumor nodularity are associated with volumetric response during adjuvant TMZ in GBM patients receiving standard of care treatment. Most notably, we found that nodular tumors have a cycle-dependent and more favorable image-based response to TMZ compared to diffuse tumors. While MGMT methylation is often considered to predict a positive response to TMZ, our results suggest that nodularity may also serve as a predictor of response, especially among unmethylated tumors.

## Author contributions

Conception and design: SCM, HW, KRS; Querying and development of cohort: HW, SKJ, AHD, KS; Acquisition of data: SKJ, PRJ, MMM, BRB, ABP, SV, JNS; Analysis and interpretation of data: PW, SCM, KRS; Writing of the manuscript: PW, TD, SCM; Review and/or revision of the manuscript: All authors.

