## Supplement for "Image-based metric of invasiveness predicts response to adjuvant temozolomide for primary glioblastoma"

### Supplement 1: Nadir

| | N= | Mean | Median | Range |
|---|---|---|---|---|
| Radii at Nadir (T1Gd) (mm) | 90 | 10.34 | 9.645 | 0-22.22 |
| %ΔT1Gd-Nadir (From pre-adjuvant to nadir) | 90 | -11.19 | -11.86 | -100 - 215.7 |
| Time to nadir (Days from pre-adjuvant date) | 90 | 124.1 | 69 | 20-512 |
| Proportion of time through adjuvant that nadir occurred[b] | 90 | 0.7842 | 1 | 0.1386- 1 |
| Nadir timing<br>Nadir before post-adjuvant date<br>Nadir at post-adjuvant date | <br>41 (45.6%)<br>49 (54.4%) | <br>-----<br>----- | <br>-----<br>----- | <br>-----<br>----- |

Distributions and counts of variables related to nadir analysis. Nadir is defined as the lowest volume on T1Gd imaging (converted to spherically equivalent radii for analysis) after the pre-adjuvant date and on or before the post-adjuvant date.

[b]For example, a value of 1 means that nadir occurred on the post-adjuvant image date, 100% of the way through adjuvant therapy. A value of 0.5, means that nadir occurred halfway through adjuvant therapy.

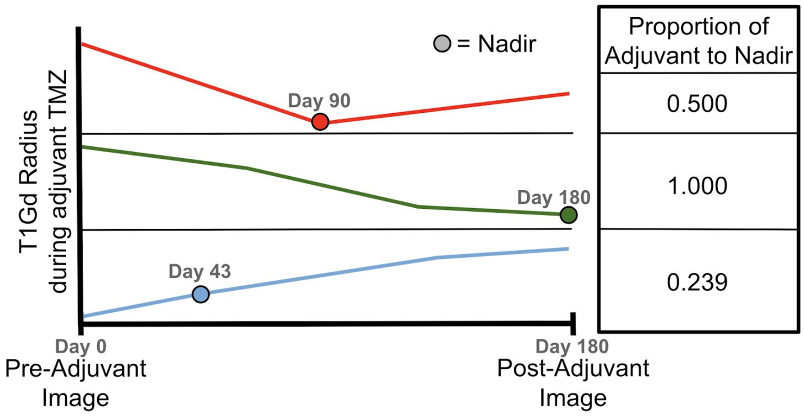


Schematic showing T1Gd radius changes from the pre-adjuvant to post-adjuvant imaging time points to demonstrate how nadir is determined and how proportion of adjuvant to nadir is calculated. Proportion of adjuvant to nadir is calculated as number of days from pre-adjuvant image to nadir image divided by number of days between pre-adjuvant and post-adjuvant imaging. Top row: Example of tumor reaching nadir volume before post-adjuvant imaging, where %ΔT1Gd-Nadir (about -0.95) is less than %ΔT1Gd (about -0.50). Middle: Example of tumor reaching nadir at post-adjuvant time point, where %ΔT1Gd-Nadir equals %ΔT1Gd. Bottom: Example of how nadir is determined when tumor volume never decreases below pre-adjuvant value.

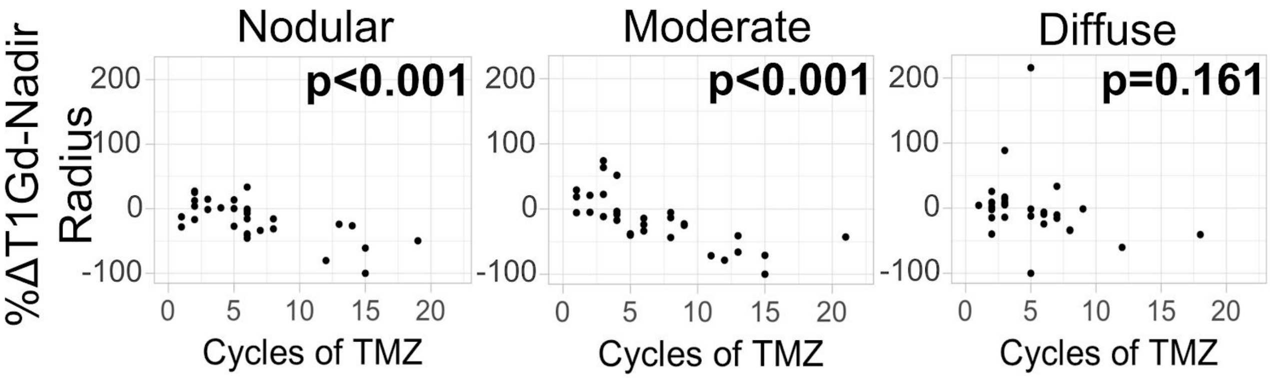


The nodular tumors have a negative correlation between number of TMZ cycles received and percent change of T1Gd radius from pre-adjuvant imaging to nadir imaging (F-test, p<0.0001), while the diffuse tumors do not have a significant trend (p=0.161), supporting the results from **Figure 4.**

**Supplement 2: MGMT Methylation Status**

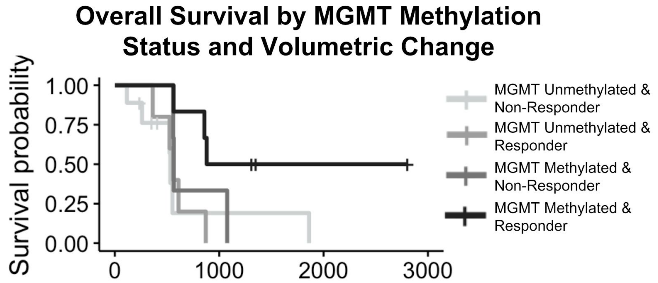


| | Total | Responder | Non-Responder |
|---|---|---|---|
| **MGMT Unmethylated** | 14 | 5 (36%) | 9 (64%) |
| **MGMT Methylated** | 9 | 6 (67%) | 3 (33%) |

Although the small sample size limits the statistical power of this comparison, methylated responders tended to have the best survival outcomes compared to other groups, followed by methylated non-responders, unmethylated responders, and unmethylated non-responders.

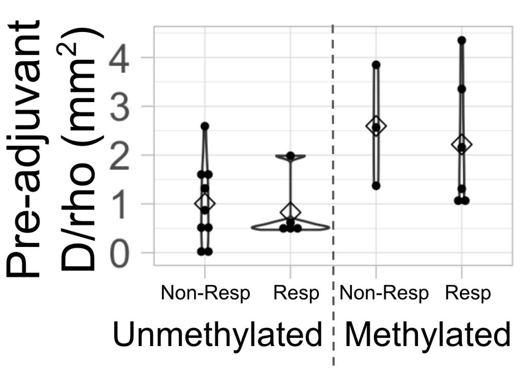


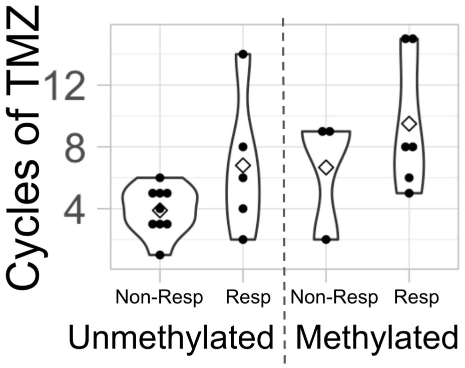


Methylated tumors are more diffuse than unmethylated tumors. Among unmethylated tumors, four of the five responders are quite nodular, while the non-responders are distributed more evenly. Methylated patients received significantly more cycles of TMZ than unmethylated patients (t-test p=0.0156). Among unmethylated patients, responders tended to receive more cycles of TMZ than non-responders.

**Supplement 3: MGMT Expanded Cohort Validation**

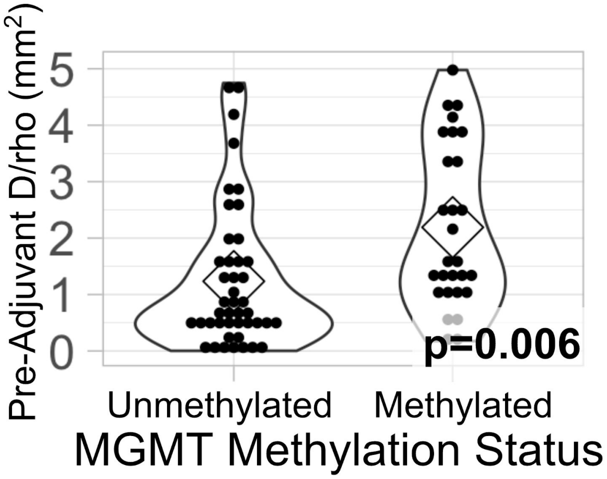


This expanded cohort (44 unmethylated, 28 methylated) uses a larger sample size to validate the observation that MGMT methylated tumors are significantly more diffuse than MGMT unmethylated tumors at the pre-adjuvant time point. Therefore, we do not think that the relationship between nodularity and response to TMZ is confounded by a predominance of MGMT methylated tumors in the nodular groups. Patients in this cohort are the 23 MGMT patients from the original patient cohort plus 49 additional patients with available MGMT status and pre-adjuvant D/rho from our database. These additional patients (30 unmethylated and 19 methylated) were first diagnosis GBM patients who received maximal safe resection followed by concurrent radiotherapy (XRT) and chemotherapy (TMZ). Pre-adjuvant d/rho was calculated from the T1Gd and T2-FLAIR images taken after concurrent therapy ended and before adjuvant therapy began. Patients who received therapies other than XRT, TMZ, steroids, or anti-seizure medications before the image was taken were not included in this cohort.

**Supplement 4: Pseudoprogression Investigation**

In order to reduce the impact of pseudoprogression on our results, we re-produced figures 1 and 3-5 using all of the patients from the original cohort that had at least 12 weeks between the end of XRT and the post-adjuvant image date (n=72).

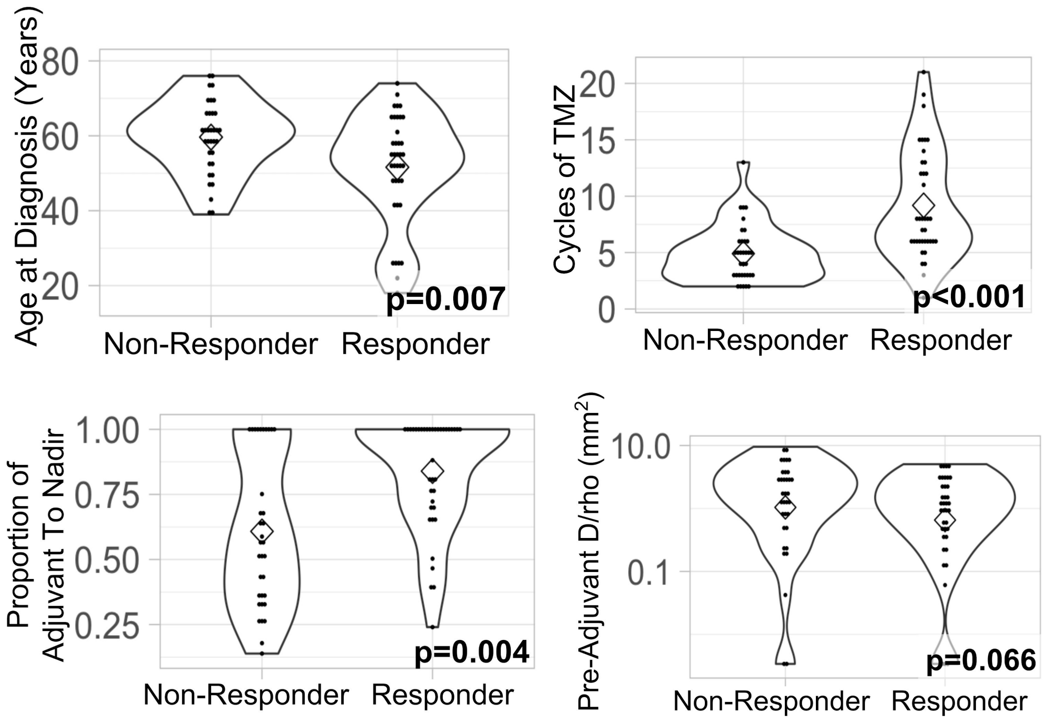


Characteristic differences between responders (n=38) and non-responders (n=34) for patients with more than 12 weeks between end of XRT and post-adjuvant imaging. These results are almost the same as those in Figure 1, except the pre-adjuvant D/rho comparison is no longer statistically significant (p=0.066).

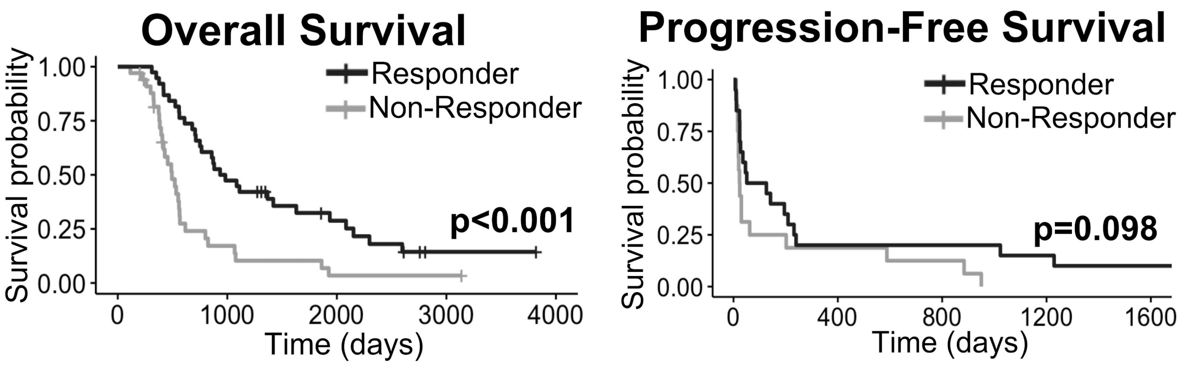


Outcome differences between responders (n=38) and non-responders (n=34), showing similar results as those in Figure 3.

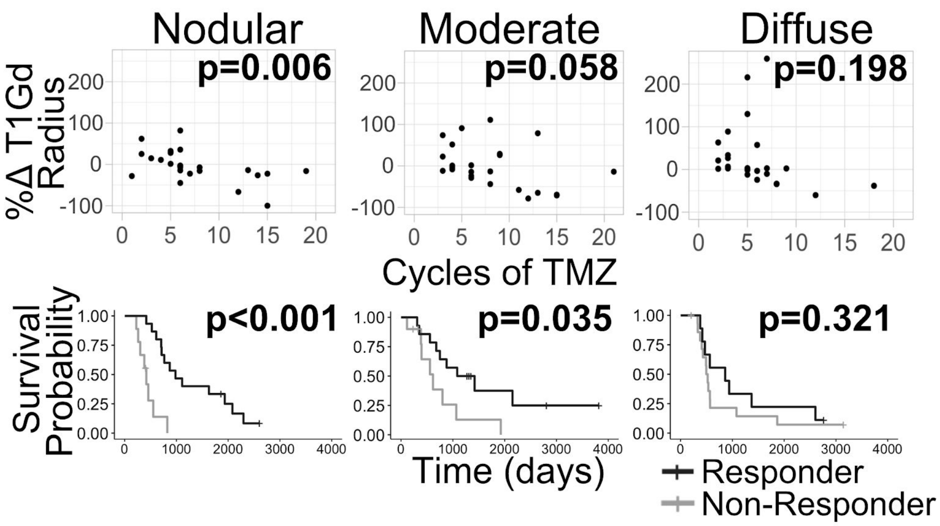


Similar to the results of Figure 4, nodular tumors (n=24) show a significant negative correlation between cycles of TMZ received and change in tumor size and this change in tumor size results in a significant survival benefit. Neither the trend nor the survival benefit are observed among diffuse tumors (n=24).

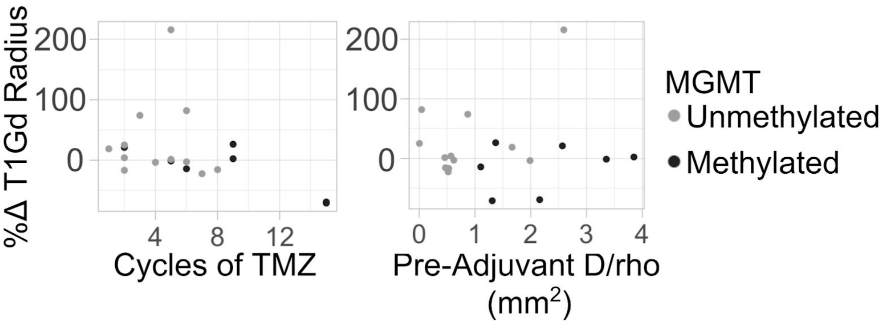


Similar to Figure 5, methylated patients (n=9) show a clearer correlation between cycles of TMZ received and change in tumor size. Unmethylated tumors (n=10). tend to be more nodular compared to the methylated ones.

**Supplement 5: Sex differences figures**

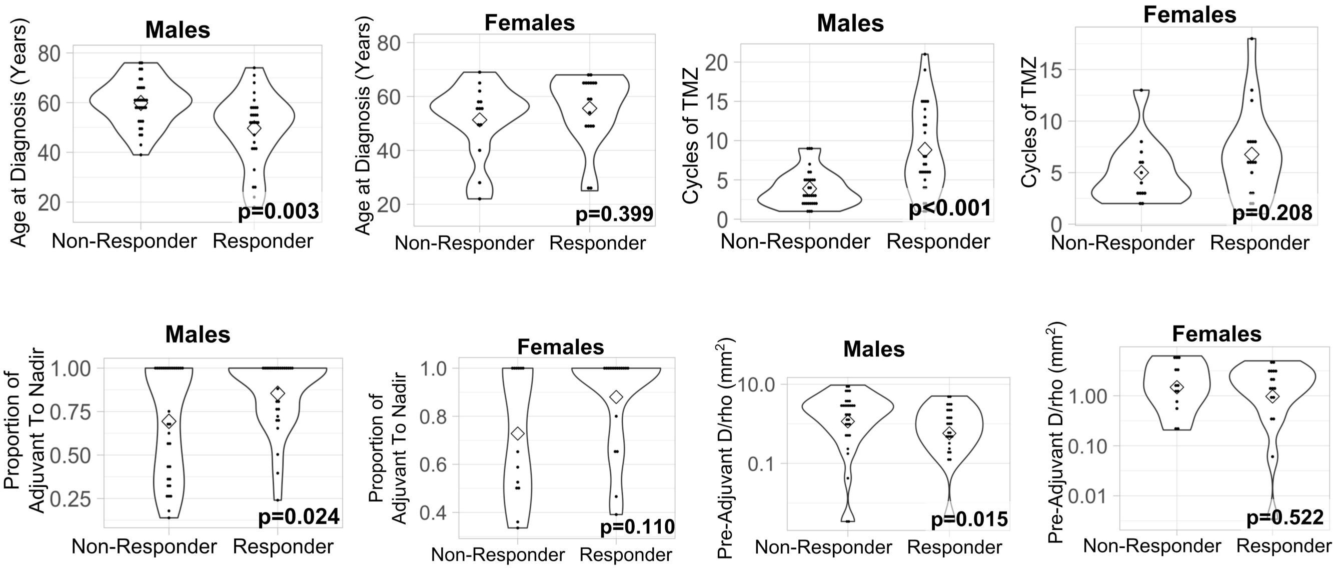


Figure 1 split into males and females. Comparison of characteristics between responders (male n=28, female n=17) and non-responders (male n=32, female n=13). Males have the same results as the combined population, while the female tests were insignificant.

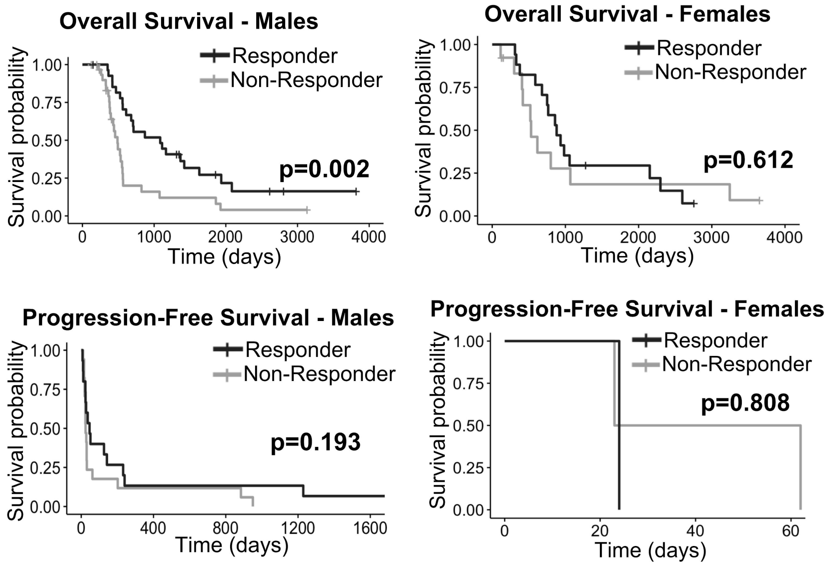


Figure 3 split into males and females. Male responders (n=28) had better overall survival than male non-responders (n=32), while females did not have a significant survival difference between responders (n=17) and non-responders (n=13). Neither males nor females had a significant difference in progression free survival between responders (male n=15, female n=1) and non-responders (male n=17, female n=2).

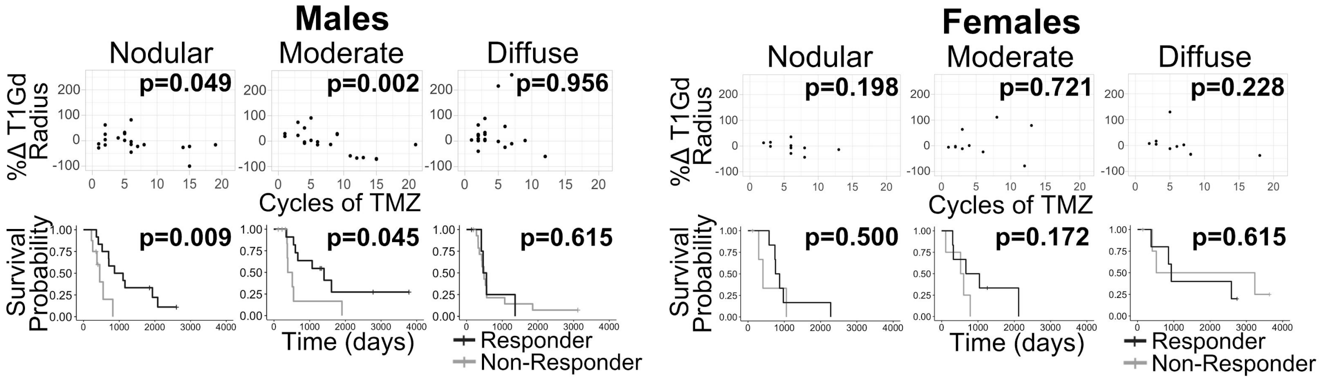


Figure 4 split into males and females. Similar to the combined population, males with nodular tumors (n=20) have a significantly negative trend between cycles of TMZ and volumetric change and this volumetric change results in a survival difference, while neither of these observations are significant among diffuse male tumors (n=20). Visually, females show similar trends as the combined population, but the small female population size (nodular n=10, diffuse n=10) likely contributes to the statistical insignificance of these observations.

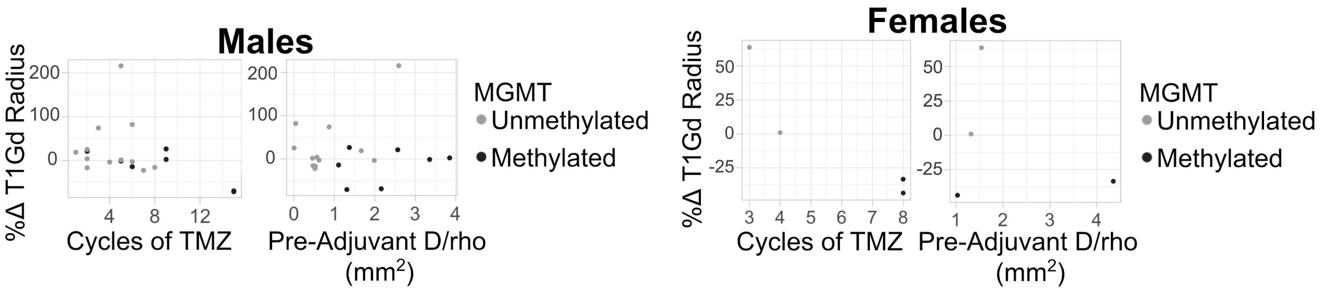


Figure 5 split into males and females. Males (unmethylated n=12, methylated n=7) show similar trends between methylation status, volumetric change, diffusivity, and cycles of TMZ as the larger population, while there are not enough females with methylation status available (unmethylated n=2, methylated n= 2) to draw a conclusion.
